# VICTOR: A visual analytics web application for comparing cluster sets

**DOI:** 10.1101/2021.03.22.436502

**Authors:** Evangelos Karatzas, Maria Gkonta, Joana Hotova, Fotis A. Baltoumas, Panagiota I. Kontou, Christopher J. Bobotsis, Pantelis G. Bagos, Georgios A. Pavlopoulos

## Abstract

Clustering is the process of grouping together different data objects based on similar properties. Clustering has applications in various case studies from several fields such as graph theory, image analysis, pattern recognition, statistics and others. Nowadays, there are numerous algorithms and tools able to generate clustering results. However, different algorithms or parameterization may result in very different clusters. This way, the user is often forced to manually filter and compare these results in order to decide which of them produce the ideal clusters. To automate this process, in this study, we present VICTOR, the first fully interactive and dependency-free visual analytics web application which allows the comparison and visualization of various clustering algorithms. VICTOR can handle multiple clustering results simultaneously and compare them using ten different metrics. Clustering results can be filtered and compared to each other with the use of interactive heatmaps, bar plots, correlation networks, sankey and circos plots. We demonstrate VICTOR’s functionality using three examples. In the first case, we compare five different algorithms on a protein-protein interaction dataset whereas in the second example, we test four different parameters of the same clustering algorithm applied on the same dataset. Finally, as a third example, we compare four different meta-analyses with hierarchically clustered differentially expressed genes found to be involved in myocardial infarction. VICTOR is available at http://bib.fleming.gr:3838/VICTOR.

## INTRODUCTION

Clustering is the process during which objects with similar properties or features are assigned to the same cluster, whereas objects with dissimilar properties are assigned to different clusters. Clustering is applicable to a wide range of scientific fields varying from Biology [1], Medicine [2], and Computer Science [3], to Physics, Chemistry and Marketing research [4]. In the Biomedical field for example, hierarchical clustering can be used to cluster data on a heatmap, infer phylogenies or group data based on similarities (e.g. sequence similarities or gene expression patterns). Another application of clustering is the detection of communities or densely connected regions in networks [5]. An example could be the detection of strongly connected components (e.g. proteins or genes in a protein-protein interaction (PPI) network), which might be involved in similar biological processes or pathways. In single cell RNA sequencing (scRNAseq) experiments, clustering based on dimensionality reduction methods can be used to group together cells with similar gene expression profiles. In Medicine, signal processing based algorithms can be used to identify objects in an image and act as automated diagnostic tools. In the field of metagenomics, binning can be used to group contigs and assign them to individual genomes [6].

Clustering algorithms generally fall into one of the four main categories based on the adopted method for defining groups. These are: hierarchical, partitional, density-based and grid-based [7], [8]. Hierarchical clustering normally uses a tree representation to visualize clusters. Widely used agglomerative or divisive hierarchical clustering algorithms include the single-, centroid-, average- and complete-linkage as well as Neighbor Joining or UPGMA [9]. Partitional algorithms assign objects into non-overlapping clusters by trying to minimize an objective function, like the summation of distances from designated centroids. K-means [10], [11], PAM [12], CLARA [13] and CLARANS [14] are typical representatives of this category. Density-based clustering is a methodology that forms clusters by identifying high-density regions. Clusters are separated by sparser data areas whose points are considered noise, borders or outliers. Two of the mostly used algorithms in this category are DBSCAN [15] and DBCLASD [16]. Grid-based clustering algorithms are applied on multi-dimensional data and divide space into cells, forming a grid. The data density in various segments of the grid indicates whether a cluster is going to be formed or not. STING [17] and CLIQUE [18] algorithms are typical examples in this category. As far as network clustering is concerned, graph-based clustering analysis can exploit the topology of a network for community detection [19]. Widely used graph-based clustering algorithms include Louvain [20], SPICi [21], Markov clustering (MCL [22]/HipMCL [23]) and Walktrap [24].

Several implementations of the aforementioned algorithms can be found in established platforms. The Weka data mining software [25] for example, supports clustering methods such as expectation-maximization [26], filtered clustering [27], hierarchical clustering [28] and k-means [29]. ClusterMaker [30], a Cytoscape [31] plugin offers both network and feature clustering. Network clustering algorithms include MCL, Affinity Propagation [32], MCODE molecular complex detection (MCODE) [33], community clustering [34], spectral clustering of protein sequences [35], TransClust [36], and AutoSOME [37]. Attribute clustering algorithms include hierarchical, k-means, k-medoid [38] and AutoSOME. NORMA [19], a web server visualization application focusing on the visualization of network annotations, is supported by the igraph library [39], thus allowing the utilization of clustering algorithms such as fast-greedy, Louvain and label-propagation [40].

Despite the plethora of clustering algorithms, when applying them to datasets, results may differ significantly upon parameterization or algorithm selection. In addition, in most cases, predefined groupings which can be used for benchmarking are rarely used. Therefore, it is often difficult to argue which algorithm or parameterization works best for certain use cases. Therefore, various studies have tried to describe protocols on how to compare clustering results, benchmark them and evaluate clustering schemas [41], [42]. Such implementations to serve this purpose already exist. XCluSim [43] for example is a standalone Java application for data clustering and visual clustering comparison and is only available upon request. XCluSim allows users to either run clustering algorithms from the Weka package within the application or upload their own external clustering results and directly compare them. However, most of the comparisons are measured using only the F-Measure (F1) metric [44]. XCluSim provides three main overviews for visualizing cluster sets comparisons. These are: *i*) a force-directed network with pie charts (object participation in clusters), *ii*) a dendrogram view and *iii*) several parallel views (e.g. Sankey plots) for more in-depth comparisons. Moreover, MatchMaker [45] is a Java implementation for visualizing cluster sets comparisons and is offered as part of the Caleydo visualization framework [46]. MatchMaker generates a combination of heat maps and parallel view plots. There are two Windows applications relative to cluster sets comparison visualization. These are: *i*) HCE [47] for hierarchical heatmaps/dendrograms and *ii*) CComViz [48] for parallel set viewing.

To go beyond current implementations, in this article, we present VICTOR, the first visual analytics web server application which offers interactive comparisons of clustering results using a plethora of metrics. Users can choose among multiple interactive visualizations such as bar plots, histograms, networks, sankey plots, circos plots and hierarchical heatmaps. VICTOR is of general purpose and fully independent as it does not come with integrated clustering algorithm implementations like in other applications. It is written in R, Shiny and JavaScript and its backend calculations are mainly based on the mClustComp library [49]. Overall, we expect VICTOR to be applicable to many scientific areas and become the reference tool for automated and sophisticated cluster sets comparison and analytics.

## MATERIAL AND METHODS

### Metrics for comparing cluster sets

In its current version, VICTOR supports ten metrics for comparing cluster sets, offered by the mClustComp library [49]. As a cluster set we define the result of a clustering analysis procedure. A cluster set consists of clusters containing objects/elements. These metrics are grouped into three main categories; *i*) Counting Pairs, *ii*) Set Overlaps/Matching and *iii*) Mutual Information [50], [51]. Metrics in the first two categories exploit the confusion matrix between cluster sets while metrics of the third category utilize the entropy of information. The ten metrics which are supported by VICTOR are: *1*) Adjusted Rand Index, *2*) Fowlkes-Mallows Index, *3*) Jaccard Index, *4*) Overlap coefficient, *5*) Wallace criteria type 1, *6*) Wallace criteria type 2, *7*) Maximum-Match Measure, *8*) Normalized Mutual information by Strehl and Ghosh, *9*) Normalized Mutual information by Fred and Join and, *10*) Normalized Variation of Information.

#### Confusion matrix

Let *C* = {*C*_1_, *C*_2_, …, *C*_*k*_} be a cluster set composed by *k* non-empty clusters of a dataset *D* and *C*’ = {*C*_1_, *C*_2_, …, *C*_*l*_} a second cluster set consisting of *l* non-empty clusters of the same dataset. The *k* × *l* confusion matrix between sets *C* and *C*’ is defined as *M*_*ij*_ with each *m* element denoting the number of common elements (intersection) between the clusters *C*_*i*_ and *C*’_*j*_.

##### Category 1. Counting Pairs

In this category, calculations are based on counting pairs of co-clustered (or not co-clustered) objects in *C* and *C*’. Each pair of objects participates into exactly one of the following four categories:

*n*_*00*_ : total object pairs that are co-clustered in both *C* and *C*’.

*n*_*0*1_ : total object pairs that are co-clustered in *C* but not in *C*’.

*n*_1*0*_ : total object pairs that are co-clustered in *C*’ but not in *C*.

*n*_11_ : total object pairs that are not co-clustered neither in *C* nor in *C*’.

All of the aforementioned calculations obey the rule: 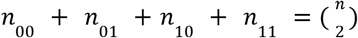.

The ***Rand Index*** measures the level of agreement between *C* and *C*’ as the fraction of agreeing object pairs to all possible pairs. It is calculated as 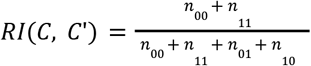 and ranges from 0 to 1 [52]. In the study of Fowlkes and Mallows [53], it is shown that for a large number of clusters relative to the total number of objects, the Rand Index undesirably converges to 1. For this reason, in VICTOR, we use the Adjusted Rand Index as an alternative.

The ***Adjusted Rand Index*** is an adjustment of the Rand Index proposed by Hubert and Araabie [54] and is defined as: 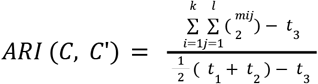 where: 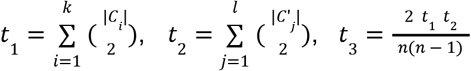[51]. Intuitively, the Adjusted Rand Index describes the normalized difference of the Rand Index and its expected value under a null hypothesis. Specifically, this null hypothesis describes independent cluster sets based on the assumption that the confusion matrix is constructed from the hypergeometric distribution, where objects are assigned randomly in the original number of clusters (per cluster set). The Adjusted Rand Index ranges from [-1, 1], where 1 means identical cluster sets while values in the range [-1, 0] indicate independent sets [51], [54].

The ***Jaccard Index*** is equal to the number of co-clustered objects in *C, C*’, divided by the total number of object pairs excluding those that are not co-clustered either in *C* or in *C*’(*n*_11_). It is defined as 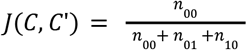 [51] and its values range from[0, 1]. Higher values indicate higher agreement between cluster sets.

The ***Overlap Coefficient*** measure is defined as the size of the intersection between two cluster sets divided by the smallest size of the two sets. The Overlap Coefficient takes ranges from [0, 1] and is calculated as 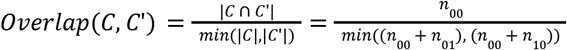 [49], [55].

The ***Wallace Criteria Type I & Wallace Criteria Type II*** are asymmetric measures that represent the probability of object pairs which are co-clustered in *C* to also be co-clustered in *C*’ for criterion type I and vice versa for criterion type II. These criteria are calculated as 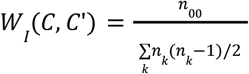 and 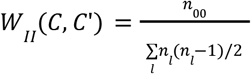 [50], [56] where *n*_*k*_ and *n*_*l*_ are the cluster sizes.

The ***Fowlkes–Mallows Index*** is symmetric and is defined as the geometric mean of the Wallace Criteria *W*_*I*_ and *W*_*II*_. It is calculated as 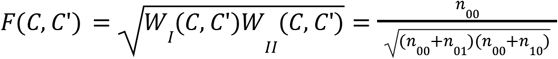[50], [51], [53] and ranges from [0, 1]. The measure is directly proportional to the number of true-positive values and therefore, a higher index indicates greater similarity between two cluster sets (1 denotes identical sets).

##### Category 2. Set Overlaps/Matching

Metrics based on set overlaps map clusters among cluster sets based on their maximum object overlap. In VICTOR we offer one metric from this category, namely ***Maximum-Match Measure***. In ***Maximum-Match Measure***, each cluster in *C* is recursively assigned a “best match” in *C*’. The largest entry of the confusion matrix *max*(*M*) is considered a maximum match between clusters *C*_*a*_ and *C*’_*b*_. The *a*-th row and the *b*-th column of the confusion matrix are recursively matched and then deleted until the matrix is empty. The Maximum-Match Measure equals to the sum of the matches divided by the total number of elements and is calculated as 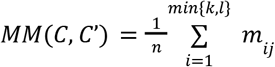 [51] where *n* is the total number of elements, *k* and *l* are the respective sizes for cluster sets *C* and *C*’and the measure values range from [0, 1].

##### Category 3. Mutual Information

All metrics in this category are based on the entropy of information and the probability of an element to be in a particular cluster. The entropy of a cluster set measures the uncertainty about the cluster in which a random object may belong to and is calculated as 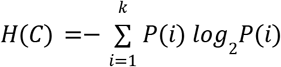 [5], [51]. *k* is the total number of clusters in cluster set *C* and 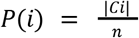 shows the probability a randomly picked element to belong to cluster *C*_*i*_ *ϵ C*. Intuitively, the ***Mutual Information*** describes how much we can reduce the uncertainty about the cluster of a random element when knowing its cluster in another cluster set (with the same set of elements). It is calculated as 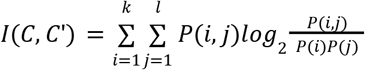 [51] where 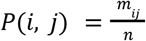is the probability of an object to belong to cluster *C*_*i*_ in *C* and to cluster *C*’_*j*_ *in C*’. Mutual Information is not bounded by a constant value which makes it more difficult to interpret. Therefore, in VICTOR, we use its normalized versions.

The ***Normalized Mutual Information by Strehl and Ghosh*** [57] normalizes the Mutual Information metric by the geometric mean of the entropies of the individual cluster sets. It is defined as 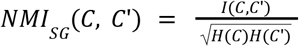 and ranges from [0, 1] with *NMI*_*SG*_ (*C, C*’) = 1 for *C* = *C*’ and *NMI*_*SG*_ (*C, C*’) = *0* if all *P*(*i, j*) = *0*or *P*(*i, j*) = *P*(*i*)*P*(*j*).

The ***Normalized Mutual Information by Fred and Jain*** [58] is another normalized version of the Mutual Information which accounts for information from both cluster sets and their total entropies. It is calculated as: 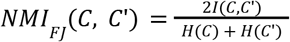 and ranges from [0, 1] like previously.

The ***Variation of Information*** metric describes the amount of information that needs to be added when going from cluster set *C* to *C*’, as well as the amount of information that is lost by *C* [50]. It is calculated as: *VI* (*C, C*’) = *H*(*C*) *+ H* (*C*’) *–* 2*I* (*C, C*’). Similarly to the simple Mutual Information measure, the Variation of Information is not bounded by a constant value and therefore, VICTOR uses its normalized version.

The ***Normalized Variation of Information*** [59] measures the average relative lack of information to infer *C* given *C*’ and vice versa. The metric is given by the formula: 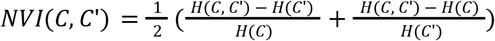. *H*(*C, C*’) is the joint entropy. Normalized Variation of Information ranges from [0, 1] and specifically the value 0 describes two identical cluster sets. In VICTOR we inverse this metric by subtracting all its values from 1, so the color coding of the various views comply with the rest of the metrics.

### Input files and dynamic filtering

In its current version, VICTOR accepts multiple clustering result (cluster set) files as input in a text format. A cluster set file normally consists of two columns without headers. The first column shows the cluster names (one cluster per row), whereas the second column shows the cluster members in comma-separated format. Users can upload multiple cluster set files and handle them simultaneously by selecting, deleting, replacing or renaming them at any time point during VICTOR’s execution. Once uploading has been completed, basic statistics like the number of clusters and number of objects are visualized as bar charts whereas histograms show an object’s participation in the various clusters.

Due to the properties of the underlying mClustComp cluster-set comparison library, each object must belong to only one cluster (no duplicates allowed). In addition, all input cluster sets which will be compared must contain exactly the same elements. This is visually shown in the ‘*Compare Clusterings*’ tab where users can observe whether all cluster sets are composed by the same number of elements, in a bar chart (Figure 1A). In case the selected cluster set files are of the same size (in terms of participating objects), users may select any of the offered metrics and proceed with comparing cluster sets.

**Figure 1.**
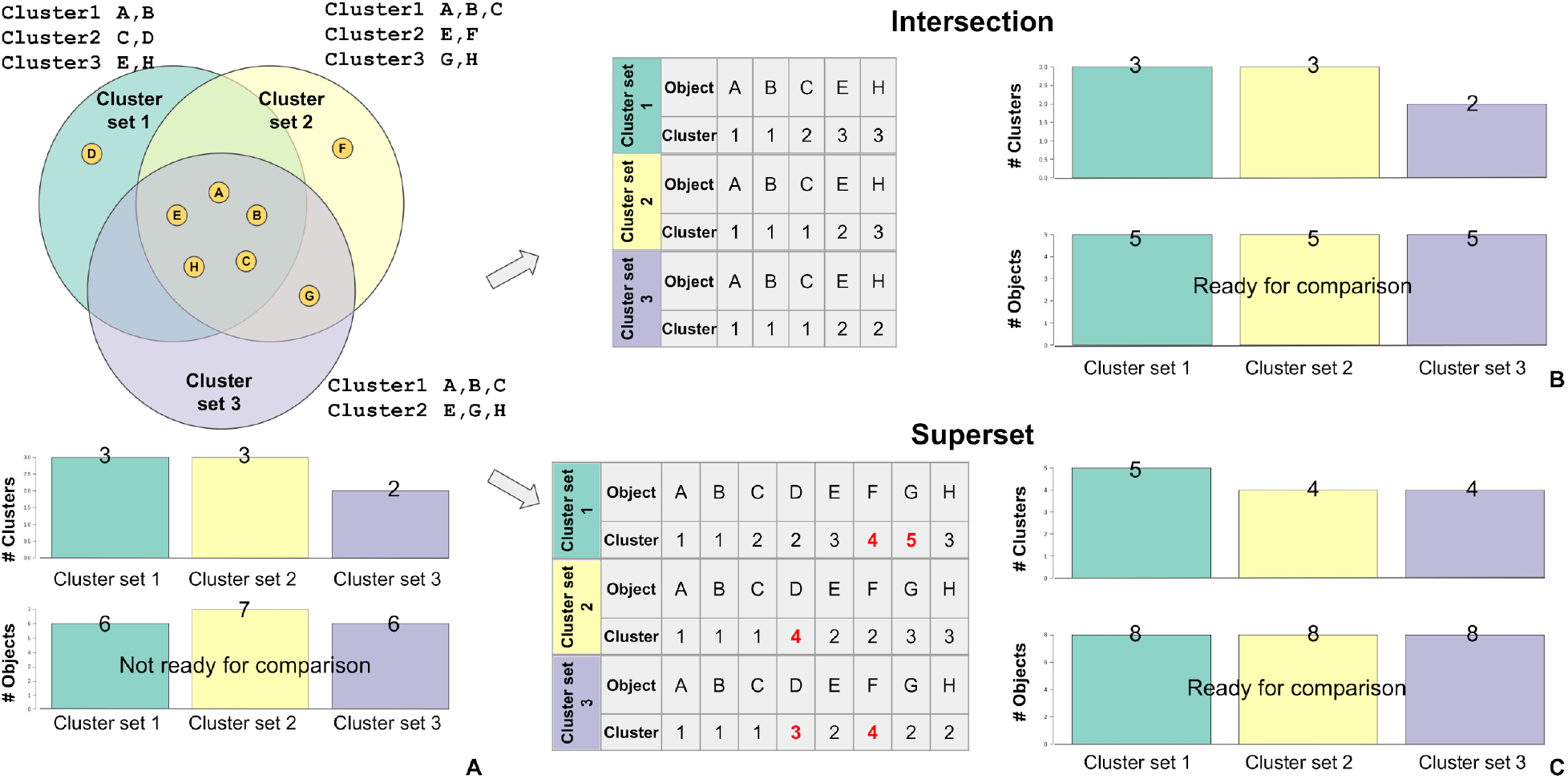
Filtering options. A) *Up:* Three cluster sets of a sample dataset consisting of 8 total objects. *Down:* The numbers of objects in the cluster sets are uneven and therefore cluster set 2 cannot be compared to the other two. B) The intersection filtering option retains only objects that are common among all selected cluster sets. C) For every object that is present in a cluster set but is missing from the others, the superset filtering option constructs a cluster represented by a singleton and assigns it to them. For example, objects F and G are missing from cluster set 1, so singleton clusters 4 and 5 are added in cluster set 1.

In the case where this does not happen, users can apply filters offered by the *File Handling* tab. Users can initially apply thresholds on the number of participating objects per cluster and keep the clusters with sizes below and above certain values. This way, users can manually eliminate singletons or exclude very large and ambiguous clusters.

Another two options which can be applied on the multiple cluster sets of interest, are to either keep the elements from the intersection of the selected cluster sets or to form a superset by adding a cluster represented by a singleton in each of the clustering results which do not contain this object (Figure 1). According to the mClustComp library, the vectors which represent the cluster sets must have the same size and be co-sorted according to the underlying objects in a way that a cluster label of an object from one cluster set (vector) is placed in the same position in the other vector. In conclusion, the intersection option maintains objects that already exist across all selected cluster set files (Figure 1B) whereas the superset option constructs singleton clusters for any of the missing objects (Figure 1C).

### Visual comparisons

VICTOR comes with a plethora of options to visualize comparisons among different cluster sets. Users may select several cluster set files of interest and any of the ten calculated clustering comparison metrics. A slider allows the interactive adjustment of the value range of the user-selected metric, which can be utilized to apply an upper and lower cutoff threshold on the visualized items (i.e. eliminating weaker cluster set similarities). Afterwards, the cluster set can be visually compared with the use of *bar charts, heatmaps, sankey plots, circos plots* and *networks* as displayed in Figure 2.

**Figure 2.**
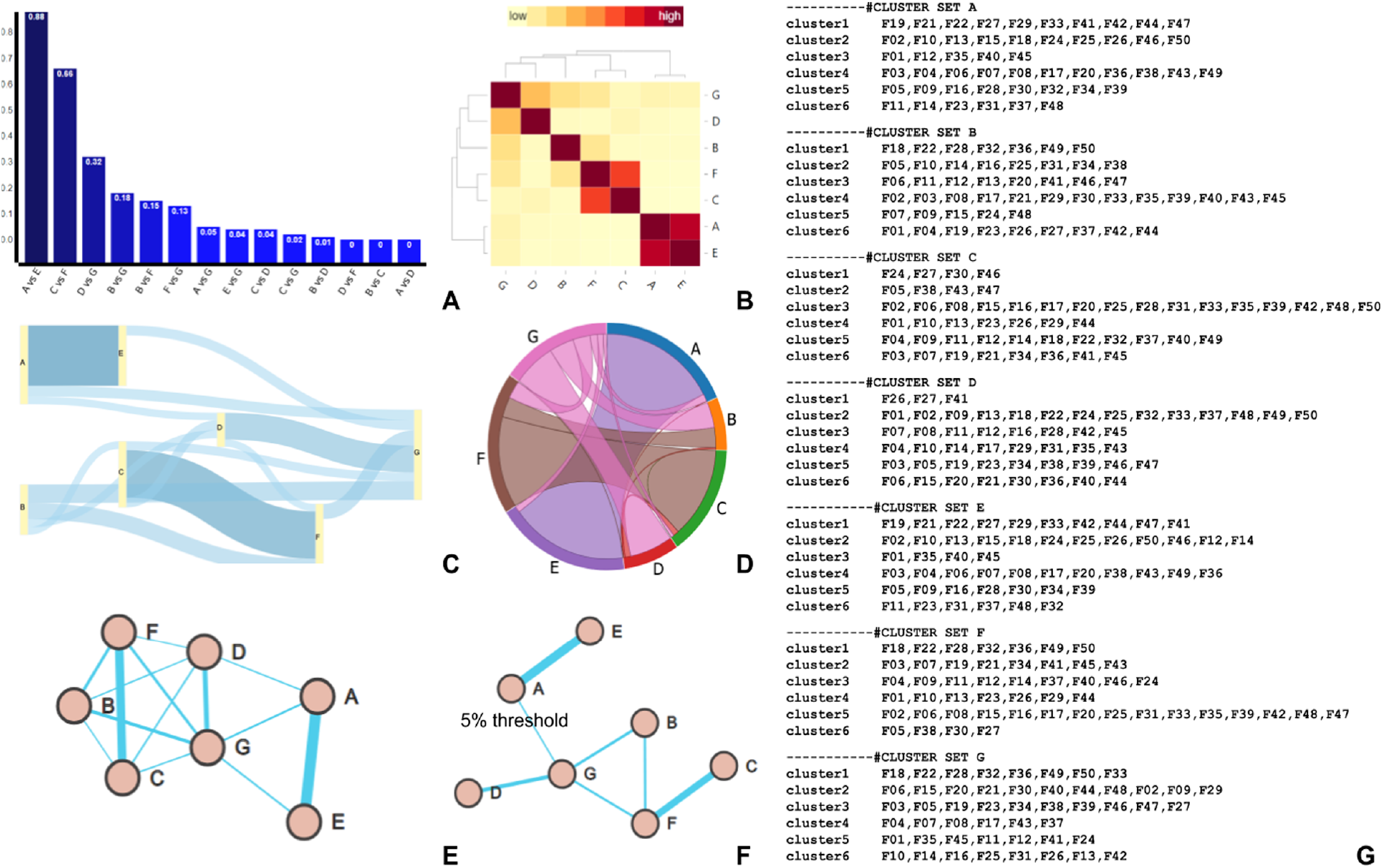
Different visualization options offered by VICTOR. Comparison of seven different cluster sets (A-F) on an artificial sample dataset. The Adjusted Rand Index metric is used for comparisons. A) Metric bar plot visualization. Each column shows the pairwise comparison of different cluster sets (A vs E, C vs F etc), with the vertical axis displaying their similarity. B) Hierarchical Heatmap. Color values range from light yellow (low similarity) to deep red (high similarity). C) Sankey Plot. Cluster sets are represented as vertical rectangles, connected by bands. The height of the band is proportional to the similarity between cluster sets; for example schemes A and E are highly similar. D) A Circos Plot visualization. Each cluster set is represented using a different color. The similarities between different schemes are represented by ribbons connecting them. E) Network visualization, after applying a force directed layout. Cluster sets are shown as nodes connected by edges, with the edge width (interaction strength) being proportional to the similarity between the cluster sets; as it can be seen, more similar schemes such as A and E, or F and C, are strongly connected, compared to other network elements. F) Network visualization after applying a 5% cut-off on the adjrand values. Application of the cut-off eliminates low-similarity connections to make the network more concise. G) The sample dataset and its cluster sets, shown in the VICTOR text input format.

Generally, Sankey plots describe flow diagrams where an arrow width is proportional to the flow rate. They are mostly used to study system energy flows [60], [61]. However, Sankey plots can also be used in biology, to plot for example risk groups against clinical cancer subtypes [62] or alternatively as a visualization option in other software suites like LiveKraken which is used for taxonomic classification [63].

In VICTOR’s sankey plots, vertical rectangles represent cluster set schemes which are then connected by undirected bands. The band height is proportional to the pairwise similarity, comparatively to the other similarity values. A heatmap is a 2D-colored representation of the value intensities of an observed phenomenon [64].

A hierarchical heatmap clusters objects in both *x* and *y* axes, according to their intensities. The color depth of a cell is proportional to the pairwise value. In VICTOR, the hierarchical heatmap is an *n x n* table, where *n* represents the number of user-selected clustering files and the color depth of each cell the value of a pairwise similarity. A dendrogram [65] is then appended on both axes, illustrating the cluster arrangement of the in-between pairwise similarities.

Bar charts [66] represent data as rectangles, where the height (or width in inverted bar plots) of a rectangle is proportional to the observed value (*y* axis). The observed data identifiers are annotated on the axis. In VICTOR, if the user does not select a cutoff threshold, 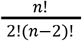 bars will be plotted, where *n* is the number of cluster sets. Each plot represents the pairwise similarity among cluster sets, based on the selected metric. Networks are also widely used to visualize data and their inter-connections in various scientific fields [5]. The observed data objects are represented as nodes and their pairwise values as edges.

In VICTOR we also visualize similarity metrics as weighted networks after applying a force directed layout algorithm such as Fruchterman-Reingold [67], Reingold-Tilford [68] and Davidson-Harel [69]. If no cutoff-threshold is selected then the drawn graph is fully connected (complete graph/clique). Subsequently, the higher the select threshold, the sparser the graph. Similarly, the stronger the similarity, the thicker the edge.

Finally, circos plots (or chord diagrams) position objects radially, where an arc length is proportional to the sum of the total interactions of an object [70]. Pairwise connections are represented as ribbons which are then colored according to one of the two interacting objects. In VICTOR, cluster sets constitute the arcs and their pairwise similarities, the ribbons. The higher the user-selected metric value threshold, the fewer ribbons will cover the circos plot.

### Conductance

VICTOR has a designated tab for calculating the conductance of a network’s clusters. In the *Conductance* tab, users can see how well-connected a cluster is to the rest of a given network, in relation to its internal connections [71]. To do that, users may upload a network file (undirected) and associate it with a clustering result of interest. The network file is normally a two- or three-column (unweighted or weighted respectively) text file in which the two first columns describe the source and target nodes and the optional third column their in-between edge weight. Notably, all objects contained in the cluster set file must be present in the network.

Let us denote a network as *G*(*V, E*) and a respective cluster set with *n* clusters. In order to calculate the conductance for each cluster, the network needs to be partitioned in two subsets *S* and *T*for each cluster *i ϵ n*. Subset *S*_*i*_ ⊆ *V*(*G*) contains all the vertices of the network that are also included in the cluster, while *T*_*i*_ ⊆ *V*(*G*) contains the vertices of the network that are not included in that cluster. The conductance for a cluster is then calculated as: 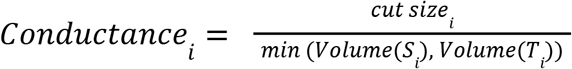 where *Volume*(*S*_*i*_) and *Volume*(*T*_*i*_) are the summation of node degrees for *S*_*i*_ and *T*_*i*_ respectively and *cut size* _*i*_ the summation of edge weights at the intersection of sets *S*_*i*_ and *T*_*i*_. The network conductance is the minimum conductance of the clusters. The closer to 0, the better the conductance of the given clusters. An example of a clustered network using two different cluster sets is shown in Figure 3. A distinct color has been assigned to each cluster and a histogram has been used to plot the conductance metric.

**Figure 3.**
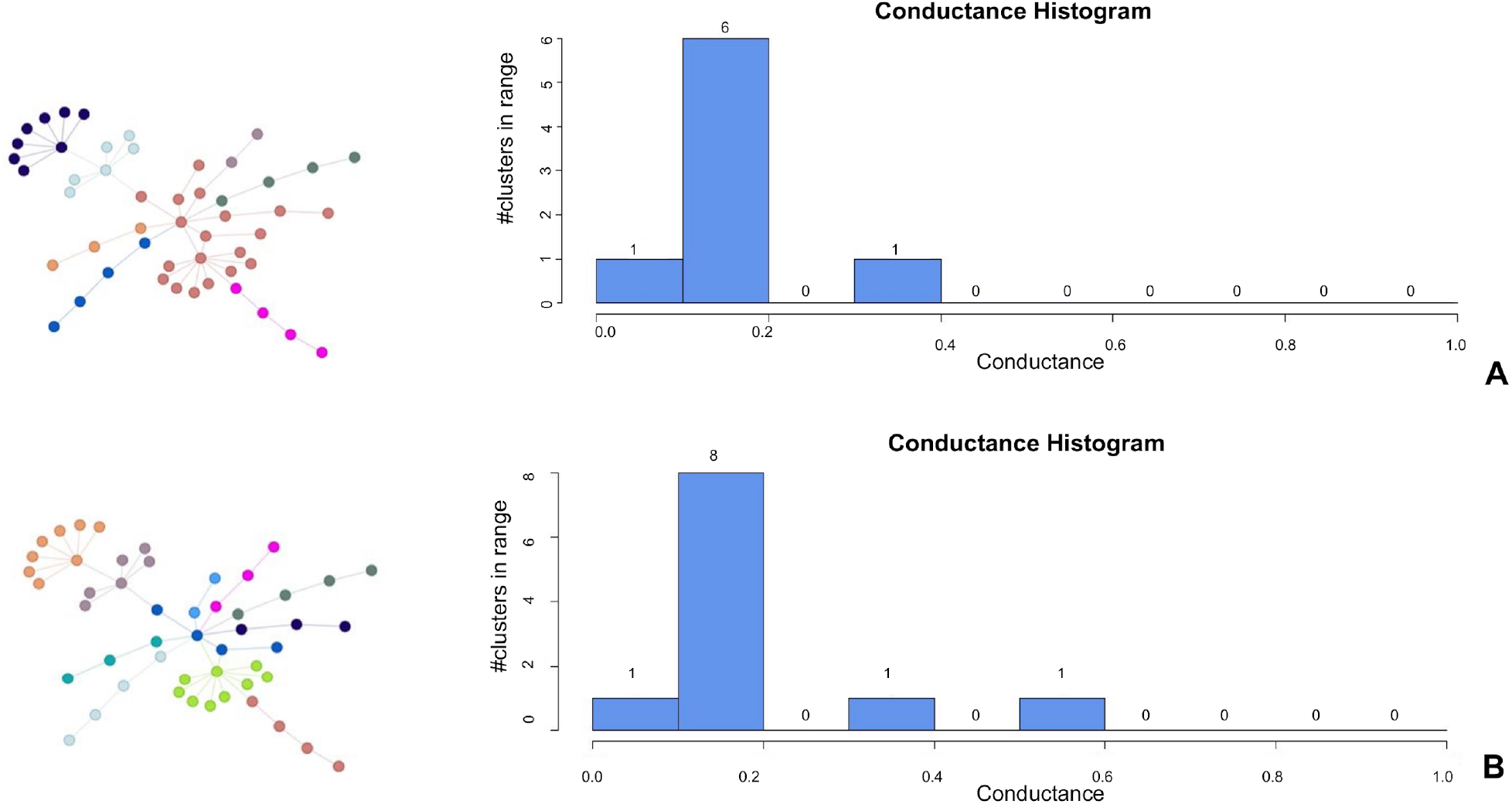
Conductance calculation for two different cluster sets by VICTOR. Conductance of two different cluster sets (A-B) on an artificial sample network. Horizontal axis of each histogram shows the conductance score whereas the vertical axis shows the number of clusters corresponding to each score. A) The clusters were generated by the Label Propagation algorithm. B) The clusters were generated by the Walktrap algorithm.

### Implementation

VICTOR is mainly written in R and JavaScript. The frontend uses the R/Shiny package, HTML, CSS and JavaScript. The Shiny package is used as the mediator to establish the communication between the R and JavaScript variables and functions. In the backend, the mClustComp library is used for the calculation of cluster-set comparison metrics. Plots are generated via the d3 JavaScript library and R.

## RESULTS

### Comparing different clustering algorithms and parameterizations

To estimate VICTOR’s efficacy, we initially tested five different network clustering algorithms applied on a Yeast protein-protein interaction (PPI) network produced using affinity purification and mass spectrometry [72]. According to NAP profiler [73], the network consists of 1,430 proteins (nodes) and 6,530 interactions (edges) and has a density of 0.01, a clustering coefficient of 0.29, and a modularity of 0.66. In order to generate clusters, we applied five clustering algorithms on the aforementioned PPI network using their default parameters. These are: *i*) MCL [74], *ii*) SPICi [21], *ii*) Louvain [20], *iv*) Walktrap [75] and *v*) Label Propagation [40].

MCL is an unsupervised clustering algorithm for graphs based on simulation of (stochastic) flow in graphs. SPICi identifies highly connected areas based on their local density. Louvain assigns nodes to communities with increased modularity. Walktrap detects communities in a graph via random walks. Finally, Label Propagation detects community structures in networks by assigning unique labels to vertices and then updates them by majority voting in the neighborhood of the node.

MCL with an inflation value of 2.0 reported 70 clusters in total. 66 of them had more than one member (corresponding to 1,426 nodes) whereas 4 nodes remained as singletons. Similarly, the SPICi algorithm (*T*_*density*_ and *T*_*support*_ set to 0.3) reported 148 clusters. 107 of them had more than three members (corresponding to 928 nodes) and 41 of them had exactly three members (corresponding to 123 nodes). 379 nodes were distributed in clusters that contained less than three members.The Louvain algorithm reported 193 clusters, where 146 of them had more than one member (corresponding to 1,383 nodes) while 47 of them were singletons. The Walktrap algorithm resulted in 294 clusters. 179 of them had more than one member (corresponding to 1,315 nodes) while 115 of them were reported as singletons. Label propagation gave an amount of 249 clusters. 160 of them have more than one member (corresponding to 1,341 nodes) whereas 89 of them were singletons.

After discarding the singletons and keeping the intersection of objects, we directly compared the five different cluster sets (Figure 4A) and showed for example that the Walktrap and Label Propagation clusterings reported more similar results compared to the rest of the clusterings. In detail, according to the left heatmap (Figure 4A), the Fowlkes-Mallows Index measure gave moderate similarity, with a value of 0.408, between the Walktrap and Label Propagation clusterings. Notably, results are not catholic and may be very different when applying the algorithms on different PPIs.

**Figure 4.**
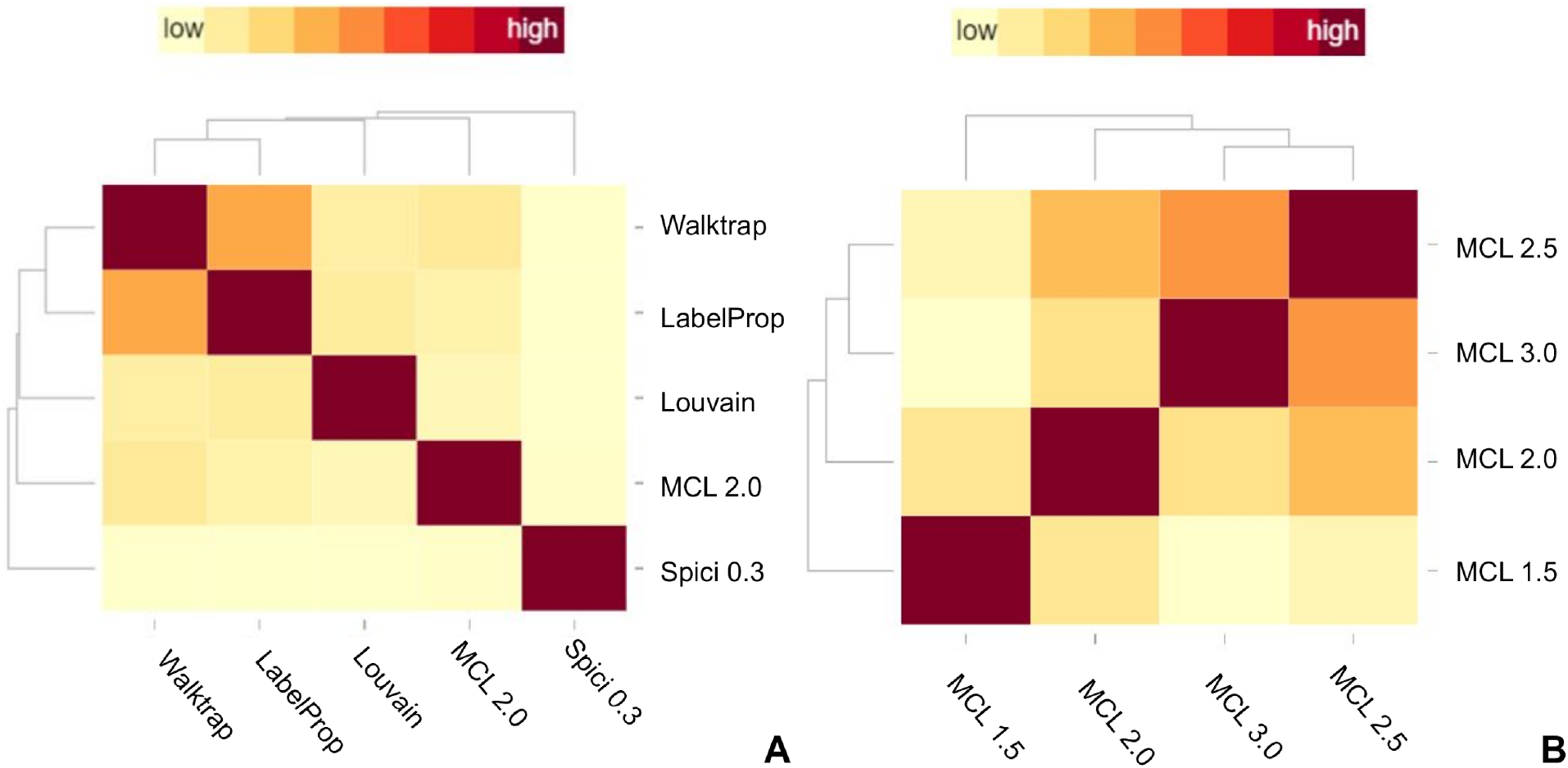
Hierarchical heatmap visualizations. A) Hierarchical heatmap of five different clustering algorithms applied on a Yeast PPI dataset. Metric used: Fowlkes-Mallows Index. B) Four different MCL clustering results for inflation parameters [1.5, 2.0, 2.5, 3.0]. The hierarchical heatmap was generated with the use of the Normalized Variation of Information metric.

To demonstrate VICTOR’s capabilities in a different scenario, we tested four different inflation parameters (1.5, 2.0, 2.5 and 3.0) for the MCL algorithm and applied it on the same Yeast PPI network. Normally in MCL, the inflation parameter adjusts the modularity and larger inflation parameters correspond to more clusters. In this run, the inflation parameter 1.5 produced 88 clusters with more than one member and 38 singletons, inflation parameter 2.0 produced 66 clusters with more than one member and 4 singletons, inflation parameter 2.5 produced 45 clusters with more than one member and 13 singletons and inflation parameter 3.0 produced 38 clusters with 37 more than two members while there were no singletons. The right heatmap (Figure 4B) was produced with the use of the Normalized Variation of Information measure and gave a moderate similarity between the inflation parameters 2.5 and 3.0 with value 0.642 and 2.0 and 2.5 with a value equal to 0.555. Like in the previous case, the results are used for demonstration purposes and may be very different on a different PPI.

### Application in gene-expression datasets from meta-analysis of myocardial infarction

For demonstration purposes, we also present the comparison of four different sets of genes from independent studies which have been used in a recently published meta-analysis for the identification of differentially expressed genes in myocardial infarction (MI) [76]. Here the setting is different: we use different (but related, since they answer the same research question) datasets and compare the clusters identified by a single algorithm. In detail, we processed four GEO microarray datasets (GSE48060 [77], GSE60993 [78], GSE61144 [78], GSE66360 [79]) and identified a total of 626 differentially expressed genes in MI patients after comparing them with healthy individuals using an FDR-adjusted p-value threshold of 0.01. The expression values for each dataset were normalized using the Quantile normalization algorithm offered by the Expander application [80] and as a next step, the genes in each dataset were clustered using an average-linkage hierarchical clustering. Four hierarchical trees were generated (one for each of the datasets) in Newick format. The trees were visualized using iTOL [81] and were used to generate clusters, by applying a distance threshold at each hierarchical tree, so that each cluster consists of at least four genes. In detail, 54 clusters were produced for the GSE48060 dataset, 52 for the GSE60993 dataset, 56 for the GSE61144 dataset and 52 for the GSE66360 dataset. These clusters are composed of 414, 603, 576 and 417 genes respectively. In Figure 5B, the Venn diagram shows the overlapping genes between the four different datasets along with the hierarchical trees produced by the average linkage hierarchical clustering. In addition, using the *Normalized Mutual Information by Strehl and Ghosh* (NMI1) comparison metric, the pairwise similarities between the clustering results of the four datasets were visualized with the use of VICTOR. As illustrated by the Venn diagram in Figure 5B, 371 of the 626 differentially expressed genes (approx. 59.26%) are common in all four datasets. 21 genes are common in three out of four sets (GSE48060, GSE60993 and GSE66360) and 205 genes are common in only two sets (GSE60993, GSE61144). At a clustering level, using the NMI1 similarity metric, the four datasets display a ∼50% pairwise similarity which complements the overlaps shown in the Venn diagram (Figures 5c, 5d). While all of the differentially expressed genes were used for this comparison (union), similar results were produced when datasets were compared based on their intersecting genes (∼48% pairwise similarity instead of ∼50%).

**Figure 5.**
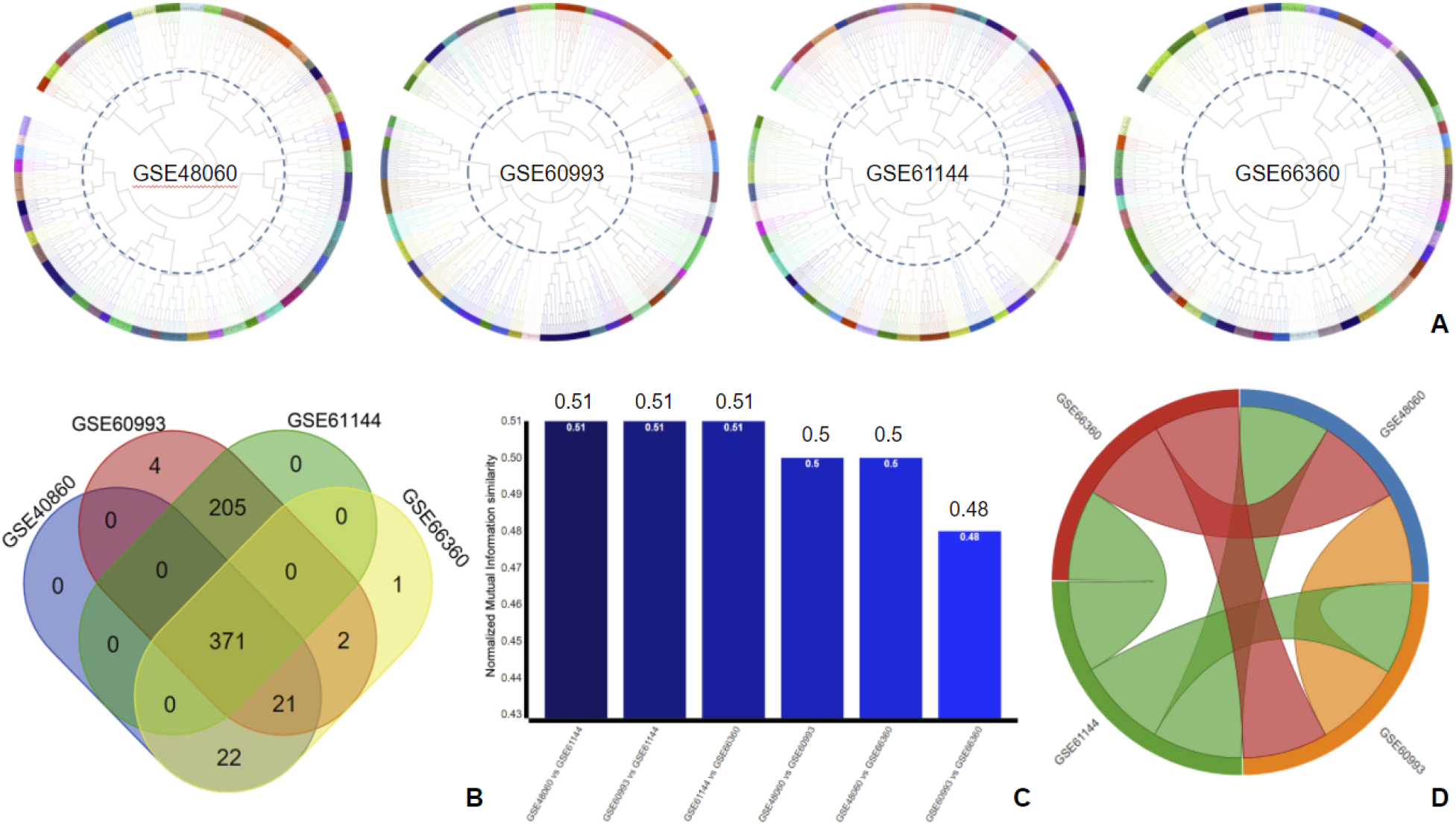
Comparison of four gene expression datasets, related to myocardial infarction. A) Hierarchical clustering of the four datasets. Clustering results are shown as circular trees. The tree leaves represent genes, which are organized into clusters and colored according to a threshold (circular dotted line). B) Gene overlaps between the four datasets are shown in a Venn diagram. C & D) Comparison of the four datasets’ clustering results using the Normalized Mutual Information (NMI) as metric. C) Pairwise NMI comparisons, shown in a bar chart. D) Circular plot showing the relationships between the four datasets.

## DISCUSSION

In this article we present VICTOR, a visual analytics tool to directly compare generated cluster sets using several comparison metrics and views. In its current version, VICTOR does not integrate implementations of clustering algorithms and therefore, it remains a flexible tool of general purpose. VICTOR’s cluster-set comparison metrics are exclusively based on the mClustComp library which returns cluster-set comparison scores using a variety of designated methods. Notably, in mClustComp, two label vectors should be of same length and this is the main reason why users are forced to choose between the intersection filtering option or the superset filtering option. In addition, in its online form, VICTOR can handle networks up to 20,000 edges and cluster set files of up to 1MB in size each, something that can be bypassed when downloading and running it locally (available on GitHub). Overall, we expect VICTOR to become a powerful tool for different communities of diverse backgrounds as well as the reference application for visually comparing cluster sets.

## AVAILABILITY

VICTOR is an open source application. Its code can be downloaded from the GitHub repository: https://github.com/PavlopoulosLab/VICTOR.

The service is available online at http://bib.fleming.gr:3838/VICTOR.

## AUTHOR CONTRIBUTIONS

EK implemented parts of the tool and wrote most of the code. MG wrote parts of the code and benchmarked the tool. IH and FAB generated the necessary data related to the first two case studies and wrote the conductance script. CB tested the tool and described the clustering comparison metrics. PGB tested the software and proposed several software optimizations and UI modifications during the development. PK provided data and the analysis for the myocardial infarction case study. GAP conceived the idea and supervised the whole project. All of the authors wrote parts of the article and have approved its final form.

